# Shotgun Metagenomic Analysis Reveals New Insights on Bacterial Community Profiles in Tempeh

**DOI:** 10.1101/2020.03.12.988444

**Authors:** Adi Yulandi, Diana Elizabeth Waturangi, Aris Tri Wahyudi, Antonius Suwanto

## Abstract

**Objective:** Amplicon sequencing targeted 16S ribosomal RNA (rRNA) has been widely used for the analysis profile of the microbial community from fermented food samples. Previous results of 16S rRNA analysis metagenome showed that *Firmicutes* was the dominant *phylum* in tempeh. However, polymerase chain reaction (PCR) steps on amplicon sequencing analysis and intragenomic heterogeneity within 16S rRNA are believed to contribute to bias in the estimation of microbial community composition. An alternative approach known as shotgun metagenomic might be able to avoid this limitation. In this study, we employed total metagenomic DNA fragments sequenced directly for taxonomic dan functional profiling analysis.

**Result:** Taxonomic profiling showed that *Proteobacteria*, *Firmicutes*, and *Bacteroidetes* were the dominant *phyla* from direct shotgun metagenomic analysis in all tempeh samples. In terms of composition, the shotgun metagenomic study revealed that *Proteobacteria* was the most relatively abundant phylum. Functional profiling showed that iron complex outer-membrane recepter protein (KEGG ID: K02014) was the most transcribed genes based on metagenome from tempeh samples.

## Introduction

Tempeh is a fermented food that originated in Indonesia. The biochemical changes of soybean during microbial fermentation increased not only nutritional values but also health-promoting bioactive compounds in the tempeh. Compared to other indigenous soybean-based fermented food such as *nato*, *miso* (Japan), *kinema* (Nepal), and *douchi* (China), which used *Bacillus* spp. as inoculum, tempeh used *Rhizopus* spp. in the production \cite{Astuti_2000}. The nature of tempeh production processes creates consortia of microorganisms not only from tempeh inoculum but also from production materials and environment. \cite{Tamang_2016}. Over the past decade, useful tools of next-generation sequencing (NGS) such as metagenomics has been applied to study microbial consortia from fermented food microbial ecology. These culture-independent techniques have two different approaches to study the taxonomic composition of the microbial community and their relative abundances. The marker-genes are amplified from total microbial genomic DNA through PCR, followed by DNA sequencing is the commonly applied option for study fermented food microbial ecology \cite{Defilippis_2017}. Other approaches called shotgun metagenomics. The total microbial genomic DNA presents in a sample will be untargeted sequencing in this approach. Besides for profile taxonomic composition study, shotgun metagenomics could be used for functional study of microbial communities based on more objective analysis through direct whole-genome sequencing \cite{Sharpton_2014, Quince_2017}. The microbial ecology study employing shotgun metagenomic analysis on samples could enhance our knowledge on both taxonomic and functional profiling of fermented food microbial community. Previous metagenomic studies microbial community during tempeh production were conducted employing amplicon sequencing targeted region V4 16S rRNA gene \cite{Radita_2017, Radita_2018, Pangastuti_2019}. These studies focused on the dynamic taxonomic profile of the microbial community from tempeh metagenome samples and indicated that Firmicutes was the predominant phylum. To enhance our knowledge on the taxonomic and functional profile of the microbial community in tempeh production, we conducted this study based on the shotgun metagenomic analysis.

## Material and Methods

### Samples

Fresh tempeh samples were collected from two local traditional producers in Bogor designated as EMP and WJB. The samples have been used as a source for microbial community analysis on tempeh for many years \cite{Radita_2018}. The EMP and WJB producer was representative for the different method in soybean boiling on tempeh production. The EMP is employing a one-time boiling while The WJB two-time.

### Total DNA Extraction

The extraction process was adapted from the previous study \cite{Seumahu_2012}. One hundred-gram of tempeh sample was homogenized in 300 mL of phosphate buffer saline (PBS) using the blender for 30 seconds. The homogenate was centrifuged at 1.000 × g for 10 minutes. Supernatants were collected and centrifuged at 10.000 × g for 3 minutes. The pellets were subjected to total microbial DNA extraction employing ZymoBIOMICS DNA/RNA Mini Kit (Zymo Research, California, USA) protocols.

### Metagenome Sequencing

The whole metagenome library preparation and sequencing process were using services from NovogeneAIT Genomics Singapore Pte Ltd. The whole microbial DNA was sheared to produce library fragments by restriction enzyme with a minimum of one μg of DNA as input. Precisely quantifying microbial DNA was used Qubit 2.0 (Thermo Fischer Scientific, United States). The purity and degradation assessment for microbial DNA was employing NanoDrop (Thermo Fischer Scientific, United States) and gel electrophoresis. The pair-end sequencing library was prepared using the TruSeq DNA PCR-Free Prep Kit (Illumina, United States). The prepared library was sequenced on the NovaSeq 6000 platform (2 × 150 bp chemistry) (Illumina, United States).

### Shotgun Metagenomic Sequence Data Analysis

The sequenced reads (raw reads) were filtered from reads; containing adapters, reads containing N (the base cannot be determined) > 10% and reads containing low quality (Qscore<= 5) base which is over 50% of the total base using NovogeneAIT Genomics Singapore Pte Ltd pipeline to produce high-quality paired-end reads (clean reads). The removal of *Rhizopus* spp reads contamination from the clean reads were employed Read QC module from MetaWrap pipeline \cite{Uritskiy_2018} using *Rhizopus microsporus* var. *microsporus* str. ATCC 52814 (GCA_002083745.1) whole-genome sequence as a reference. The SqueezeMeta pipelines \cite{Tamames_2019} were employed for assembly, taxonomic, functional and bin analysis. The pipelines used co-assembly mode option where reads from all samples are pooled before the assembly using Megahit \cite{Li_2015} step performed. The SequeezeMeta binning pipeline includes DAS Tool \cite{Sieber_2018} to merge multiple binning results from Maxbin \cite{Wu_2014} and Metabat2 \cite{Kang_2019} in just set. CheckM \cite{Parks_2015}software was used to estimate the goodness of the bins in the pipeline.

## Results

### Metagenome Sequencing and Assembly Statistic

The total microbial DNA extracted from tempeh samples collected from two different tempeh producers in Bogor, Indonesia, was subjected to Illumina whole metagenome sequencing pipelines. The average effective rate of clean reads from two raw reads metagenomic data after the quality trimming was 99.93%. Total 9,699,953 (12.28 %) reads from 78,942,348 EMP clean reads and 5,428,509 (5.02%) from 107,981,246 WJB clean reads metagenomic were mapped to the *Rhizopus* genome reference. The number of contigs results from the co-assembly steep data were 316,661. The longest contigs were 485167 bp and N50 value 1924.

### Taxonomic and Functional Profiling

The average reads it mapped with reference database using the Diamond \cite{Buchfink_2015} search on SqueezeMeta pipelines was 92.15 %. Individual gene annotation reads are used to produce a consensus for contigs. GeneBank nr was performed as a reference on taxonomic profiling. Contigs with phylum annotation were 271, 375 (85.7% from total contigs). Abundance estimation contig in each sample, the pipelines mapped clean reads onto the contigs resulting assembly step. Among bacteria, *Proteobacteria* was the abundant phylum (Figure 1). Functional profiling used the latest publicly version of KEGG database for KEGG ID annotation. Iron complex outer-membrane recepter protein (KEGG ID: K02014) was the most transcripted expression in the metagenome from tempeh samples (Figure 2).

**Figure 1.**
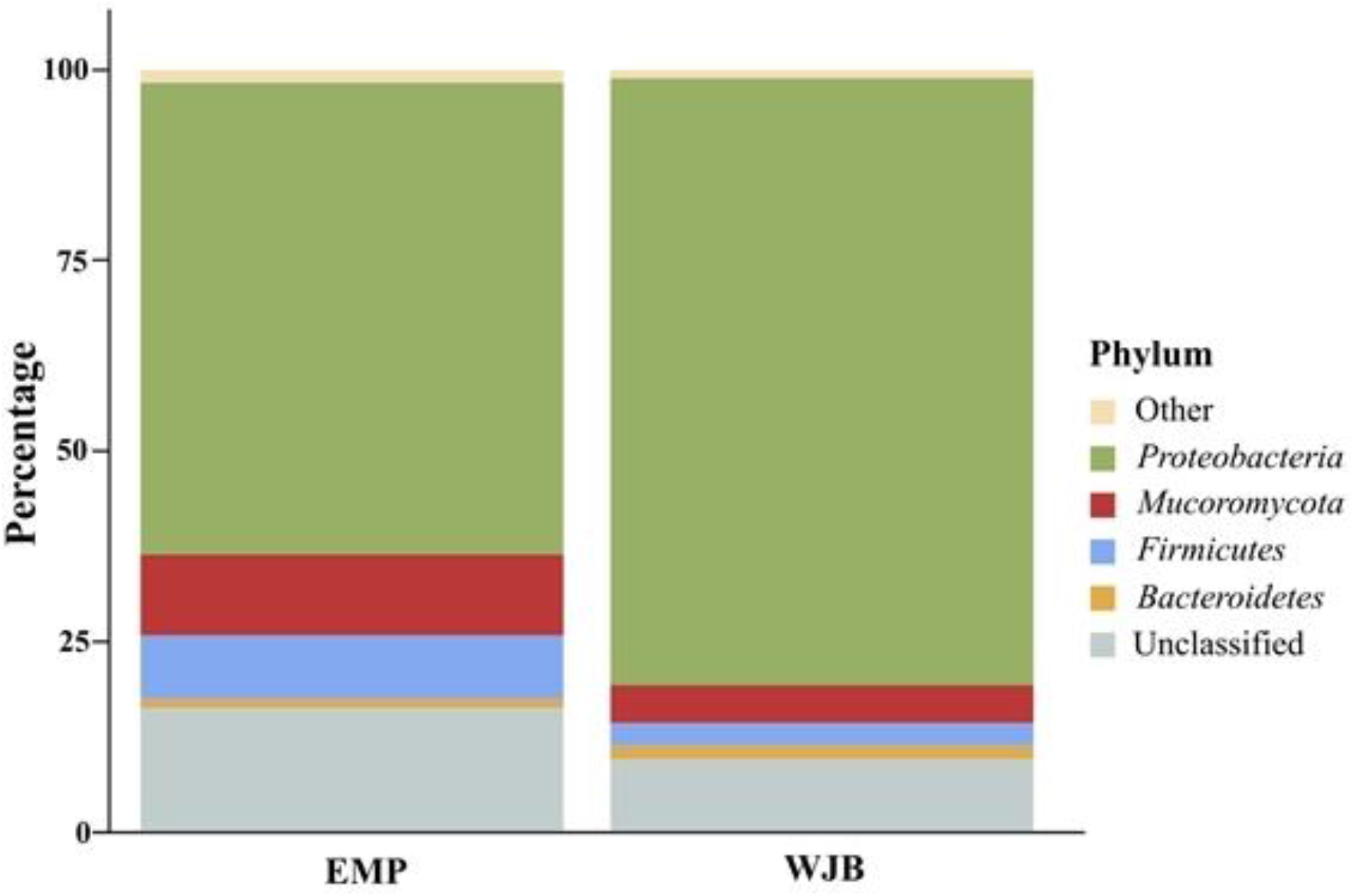
The taxonomic abundance of the microbial community tempeh samples at the rank of phylum

**Figure 2.**
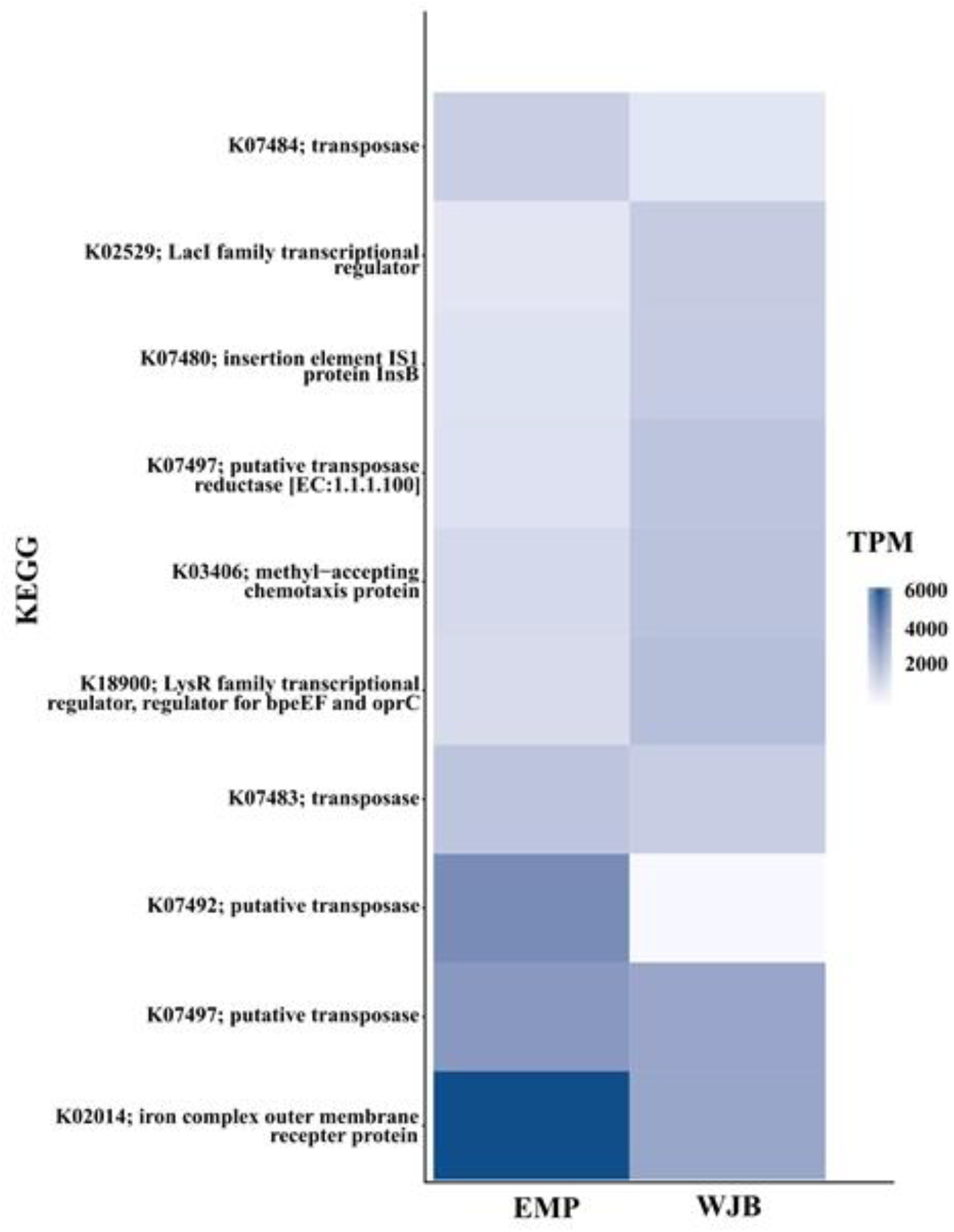
The functional profile of the metagenome tempeh samples using KEGG annotation in TPM (transcripts per kilobase million)

### Binning and Bin Check

The total number of bins obtained from co-assembly metagenome EMP and WJB samples result from DAS tool were 22. According to CheckM result, nine bins categorized as good-quality bins, which completeness more than 75% with less than 10% contamination. Among good-quality bins, six categorized as high-quality bins, which completeness more than 90% (Table 1).

**Tabel 1.**
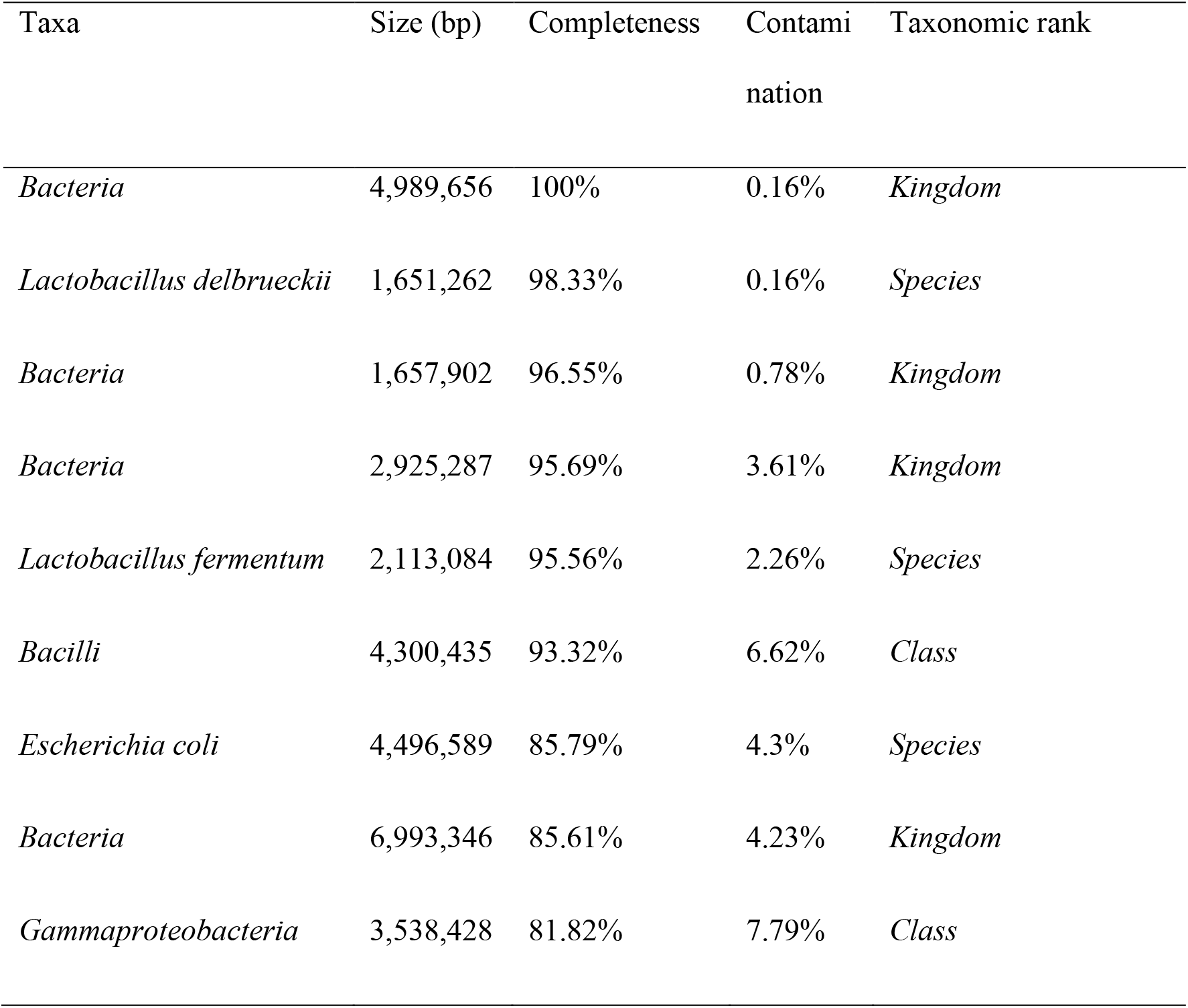
High quality bins (>90% completion, <10% contamination) obtained by co-assembly mode of EMP and WJB.

## Discussion

Study of the microbial community of EMP and WJB tempeh samples using 16S rRNA metagenome sequencing analysis already conducted. Concordant with the result of shotgun metagenome sequencing analysis in this study, *Proteobacteria, Firmicutes, and Bacteroidetes* were the abundant phylum, although different in composition. *Firmicutes* relatively most abundant in amplicon sequencing study while in a shotgun metagenomics study was *Proteobacteria*. The data expand the understanding of bioprocess during soybean fermentation compare to previous microbial ecology study using amplicon sequencing \cite{Kumar_2019, Sarkar_2002}.

## Data Availability

The tempeh metagenomic raw reads used for this study were deposit in publicly accessible NCBI’s Sequence Read Archive (SRA) under the accession number: PRJNA605305.

